# Parallel visual pathways with topographic versus non-topographic organization connect the *Drosophila* eyes to the central brain

**DOI:** 10.1101/2020.04.11.037333

**Authors:** Lorin Timaeus, Laura Geid, Gizem Sancer, Mathias F. Wernet, Thomas Hummel

## Abstract

One hallmark of the visual system is the strict retinotopic organization from the periphery towards the central brain, spanning multiple layers of synaptic integration. Recent *Drosophila* studies on the computation of distinct visual features have shown that retinotopic representation is often lost beyond the optic lobes, due to convergence of columnar neuron types onto optic glomeruli. Nevertheless, functional imaging revealed a spatially accurate representation of visual cues in the central complex (CX), raising the question how this is implemented on a circuit level. By characterizing the afferents to a specific visual glomerulus, the anterior optic tubercle (AOTU), we discovered a spatial segregation of topographic versus non-topographic projections from molecularly distinct classes of medulla projection neurons (medullo-tubercular, or MeTu neurons). Distinct classes of topographic versus non-topographic MeTus form parallel channels, terminating in separate AOTU domains. Both types then synapse onto separate matching topographic fields of tubercular-bulbar (TuBu) neurons which relay visual information towards the dendritic fields of central complex ring neurons in the bulb neuropil, where distinct bulb sectors correspond to a distinct ring domain in the ellipsoid body. Hence, peripheral topography is maintained due to stereotypic circuitry within each TuBu class, providing the structural basis for spatial representation of visual information in the central complex. Together with previous data showing rough topography of lobula projections to a different AOTU subunit, our results further highlight the AOTUs role as a prominent relay station for spatial information from the retina to the central brain.

## Introduction

Most insects rely on visual cues for accurate maneuvering, which requires appropriate processing and fast integration of various visual stimuli (Egelhaaf and Kern 2002, Heinze 2017, Mauss, Vlasits et al. 2017)). Fast decisions on whether to veer away from or approach an immobile or moving object while remaining able to quickly orientate within a complex, three-dimensional environment are key tasks for their survival (Mauss, Vlasits et al. 2017). Research focused on dissecting neural circuits in the periphery of the visual system as well as in the central brain of a large variety of insect species, more recently focusing on the genetic model organism *Drosophila melanogaster*, has provided considerable insights into how information is processed beyond photoreceptor cells (Borst 2014, Silies, Gohl et al. 2014, Behnia and Desplan 2015). Although the resolution of an insect compound eye does not rival that of a vertebrate retina (Kirschfeld 1976), neuronal elements for the internal representation of certain features of the visual world have been successfully identified: Functional studies, more recently using genetically encoded effectors in flies, have linked distinct structures of the visual system to processing discrete aspects of visual perception (Schnell, Joesch et al. 2010, Bahl, Serbe et al. 2015, Fisher, Leong et al. 2015, Ribeiro, Drews et al. 2018). Of special interest is the central complex (CX), a structure of interconnecting neuropils (named the protocerebral bridge, ellipsoid body, fan-shaped body, and noduli) located at the midline of the protocerebrum. Across insects, its various functions comprise higher locomotor control, integration of multisensory input and memory formation (Strauss 2002, Heinze and Homberg 2007, el Jundi, Pfeiffer et al. 2014, Turner-Evans and Jayaraman 2016, Varga, Kathman et al. 2017).

The CX plays an important role in processing visual information across insects, and neural pathways connecting the CX with the optic lobes have been characterized (Homberg 2015, Turner-Evans and Jayaraman 2016, El Jundi, Warrant et al. 2018, Franconville, Beron et al. 2018, Honkanen, Adden et al. 2019). In *Drosophila*, numerous studies using a variety of genetic tools described roles of the CX in visual pattern memory (Liu, Seiler et al. 2006), encoding of visual experience and self-motion (Shiozaki and Kazama 2017), flight-dependent visual responses (Weir and Dickinson 2015), sun-guided navigation (Giraldo, Leitch et al. 2018), visual landmark recognition (Seelig and Jayaraman 2015, Green, Adachi et al. 2017), including sensimotor remapping of visual information (Fisher, Lu et al. 2019), suggesting a substantial role for the CX in guiding object recognition while orientating in space. While the neuroarchitecture of the *Drosophila* CX shows clear signs of a topographic organization (Lin, Chuang et al. 2013, Franconville, Beron et al. 2018), the cellular composition and synaptic wiring diagram of neural circuits that relay spatial information from the optic lobes into the CX remain incompletely understood.

One prominent input pathway for visual information into the ellipsoid body (EB) is via distinct classes of Ring neurons (R neurons), which form a stack of several ring-shaped layers (Hanesch, Fischbach et al. 1989, Wolff, Iyer et al. 2015, Franconville, Beron et al. 2018). Afferent neurons are synaptically connected with R neurons via distinct microglomerular structures in the bulb neuropil adjacent to the EB (formerly referred to as the lateral triangle) (Ito, Shinomiya et al. 2014). These connections are distributed retinotopically, since their positions correlate to small receptive fields on the ipsilateral side (Seelig and Jayaraman 2013, Omoto, Keles et al. 2017). The transmission of spatial information from the optic lobes to the EB likely involves two synaptic neuropils: First, the R neuron dendrites in the bulb neuropil receive direct synaptic input from tubular-bulbar neurons (or TuBu neurons), originating from the anterior optic tubercle (AOTU), one of several conserved optic glomeruli (Otsuna and Ito 2006, Ito, Shinomiya et al. 2014, Panser, Tirian et al. 2016). Recent functional studies could described how R neurons inherit their receptive field properties from TuBu neurons (Shiozaki and Kazama 2017, Sun, Nern et al. 2017). Secondly, distinct classes of medulla descending neurons (medullar-tubercular neurons, or MeTu neurons) directly connect the medulla with the AOTU (Otsuna, Shinomiya et al. 2014, Omoto, Keles et al. 2017). In contrast, the majority of remaining optic glomeruli are exclusively innervated by lobula columnar (LC) neurons (Otsuna and Ito 2006, Wu, Nern et al. 2016). The AOTU is unusual in that it can be subdivided into a medially located large unit (LU; also named AOTUm (Omoto, Keles et al. 2017), receiving input from the lobula, via LC neurons), and a more lateral, small unit (SU, receiving input from the medulla, via MeTu neurons). While recent functional studies revealed that distinct visual features are encoded by different glomeruli (or their respective subunits), e.g. small objects, escape, or reaching (Wu, Nern et al. 2016, Keles and Frye 2017), spatial information should be lost in the majority of optic glomeruli, due to convergence of intermingling LC inputs (Panser, Tirian et al. 2016). However, other studies revealed that some LC afferents display some rough spatial restriction along the dorso-ventral axis of the AOTU, indicating that a topographic pathway into the central brain may exist there (Wu, Nern et al. 2016). Hence, it remains unclear whether only a rough topographic representation of visual information exists in the central brain, or whether additional, point-to-point pathways with higher resolution also exist.

Here, we show that a stereotyped topographic map is built by distinct MeTu neuron subtypes within the SU of the AOTU, which is spatially separated from LC representation in the LU. Interestingly, the overlapping dendritic fields of different MeTu subtypes in the medulla diverge into multiple parallel visual channels that are subsequently maintained via parallel synaptic pathways from the AOTU to the bulb neuropil. Within the bulb, topographic channels connect with distinct receptive fields of central complex ring neurons, whereas non-topographic channels have different R-neuron targets. Based on these data we propose a model in which specific domains of the AOTU form a central relay station for topographic information, organized in multiple parallel channels, thereby being ideally suited for visual object recognition.

## Results

### Distinct types of afferent arborizations within optic glomeruli

Optic glomeruli and olfactory glomeruli are prominent neuropil structures located in different regions of the adult brain, with olfactory glomeruli concentrated within the antennal lobes (AL) of the deutocerebrum, whereas optic glomeruli form the AOTU, the ventrolateral protocerebrum (VLP), and the posteriorlateral protocerebrum (PLP)(Fig. 1A). To determine whether a common connectivity logic is shared between olfactory and optic glomeruli, we investigated the arborization patterns of afferent fibers projecting into optic glomeruli. Olfactory glomeruli are characterized by a sensory class-specific convergence of afferent axons, each glomerulus thereby representing a unique odorant receptor identity (Laissue and Vosshall 2008) (Fig. 1B). Within each olfactory glomerulus, single sensory axon terminals arborize throughout the glomerular volume with all converging axon branches broadly overlapping and tightly intermingling (Hummel, Vasconcelos et al. 2003) (Fig. 1C).

**Figure 1:**
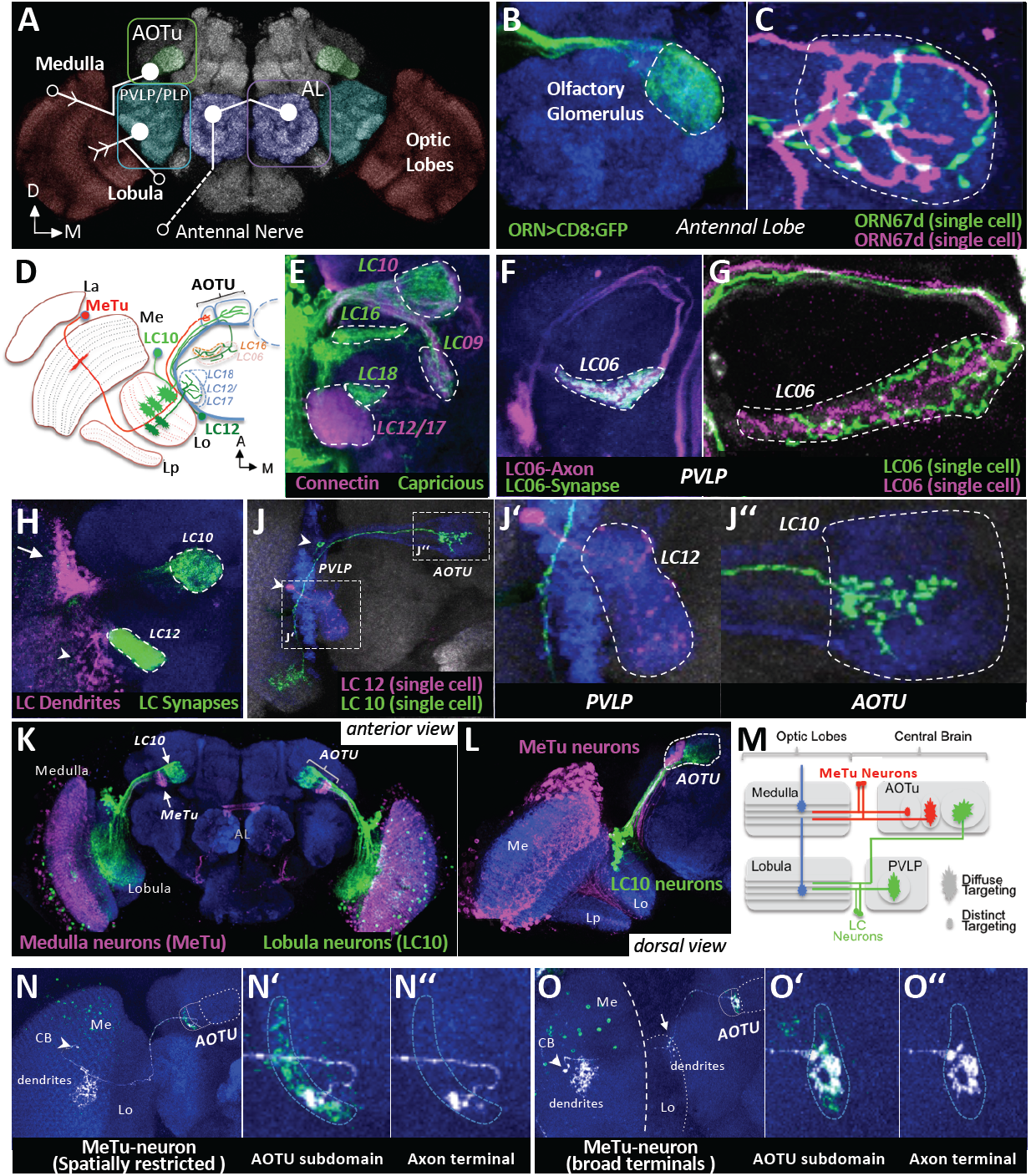
Organization of afferent projections within olfactory and optic glomeruli. **A.** Overview over sensory glomeruli. Three pathways are shown, connecting medulla, lobula and antenna with their respective target neuropils (for clarity, lobula-AOTU connections are not drawn). Open circles represent the position of the cell body, closed circles a target glomerulus and arrows indicate dendritic arborizations. AL, antenna lobes; OL, optic lobes; AOTU, anterior optic tubercle; PVLP, posterior ventrolateral protocerebrum; PLP, posterior lateral protocerebrum; **B, C.** Axon terminals of OR67d-expressing olfactory receptor neurons in the AL are branching throughout their target glomerulus DA1 and intermingle with each other. **D.** Schematic overview over visual projection neurons (VPNs) contributing to optic glomeruli (horizontal section). Only a subset of optic glomeruli are shown (the AOTU and five representatives in the PVLP). Afferents exist from the medulla (METUs cell; orange) and lobula neuropils (LC neurons; blue, green, and yellow). La, lamina; Lo, lobula; Lp, lobula plate. **E.** Optic glomeruli are marked by combinatorial expression of different cell-adhesion molecules (Connectin, magenta; Caprecious; green). **F.** LC6 terminals (marked with syt:GFP) contribute to a characteristic optic glomerulus in the PVLP. **G.** Two individual LC6 clones innervate the complete glomerulus. **H.** Co-labeld LC10 and LC12 neurons. Somato-dendritic (magenta) and presynaptic compartments (green) are labeled using DenMark and syt:GFP, respectively. Cell bodies of LC10 amarked with an arrow, LC12 with an arrowhead. **J.** Single cell morphologies of LC10 and LC12. While LC12 neurons branch throughout their target glomerulus (J’), LC10 neuron terminals are dorso-ventrally restricted within the LU (J’’). Arrowheads indicate position of cell bodies. **K, L.** AOTU compartments innervated either by METU or LC10 neurons. **M.** Schematic summary of pathways innervating AOTU and PVLP. Afferent medulla innervation indicated by blue neurons. **N-O.** Single cell clones of METU cells with spatially restricted (N) or broad axon terminals (O). Different subtypes of METU neurons can be defined based on the position and size of terminal arborizations and whether the lobula is also innervated (arrow in O). CB, cell body. For genotypes, see supplemental Data.

Inputs from LC neurons to optic glomeruli in the PLP/PLVP region are restricted to the ventrolateral brain region (Otsuna and Ito 2006) (Fig. 1D). In contrast, the more dorsally located AOTU receives afferent input via the anterior optic tract, containing both LC and MeTu fibers (Fischbach and Lyly-Hunerberg 1983, Otsuna and Ito 2006, Panser, Tirian et al. 2016, Omoto, Keles et al. 2017) (Fig. 1J). Using specific driver lines from the FlyLight and Vienna Tiles collection (Jenett, Rubin et al. 2012, Kvon, Kazmar et al. 2014), a variety of LC neuron types could be identified and their class-specific segregation into single optic glomeruli has been visualized (Costa, Manton et al. 2016, Panser, Tirian et al. 2016) (Fig. 1F-L). In analogy to work on olfactory glomeruli in the antennal lobe (Hong, Zhu et al. 2009, Hong, Mosca et al. 2012), we found that optic glomeruli can also be classified according to expression of different cell surface molecules (Fig. 1E shows an example of the expression for Connectin and Capricious in different subsets of optic glomeruli).

To characterize afferent arborizations within optic glomeruli, we generated single cell clones for different LC neuron types (LC6, LC10, LC12; Fig. 1G, H). Similar to olfactory sensory neurons (OSN) axon terminals, we found that each LC axon ramified throughout a single optic glomerulus and all neurons of the same LC class converged onto a common glomerular space (Fig. 1G, H), thereby confirming the homogeneous arborization pattern within synaptic glomeruli in the PVP/PLVP neuropil that was recently published data (Wu, Nern et al. 2016). In contrast, a more diverse pattern of afferent innervation was observed in the AOTU large and small units (Fig.1D, K). Our systematic characterization of a large collection of AOTU-specific expression lines confirmed that the LU is the target field of LC neurons whereas the SU is innervated by MeTu neurons (Fig. K-M, and see below)(Panser, Tirian et al. 2016, Omoto, Keles et al. 2017). Single LC afferent terminals in the LU arborized throughout large areas of the glomerular subunit’s volume, with some enrichment in the dorsal versus ventral regions of the LU (Fig. 1J’’)(Wu, Nern et al. 2016). In contrast, single MeTu afferents in the SU were more variable, ranging from broad (in close proximity to the LU) to spatially restricted in more lateral regions (Fig. 1N, O), indicating that different MeTu classes for distinct spatial representation might exist within the AOTU. This structural feature of non-overlapping afferent terminals is making the AOTU a candidate neuropil that maintains topographic representation of visual information within the central brain.

### Morphological and molecular domain organization of the AOTU

To determine how the architecture of the AOTU correlated with patterns of afferent innervation, we first co-labeled glial membranes with the neuropil epitope N-Cadherin (Fig. 2A, B). As previously reported (Omoto, Keles et al. 2017), a subdivision of the SU neuropil into multiple domains along the medial-lateral axis became visible, whereas the LU appears like a homogeneous neuropil without any obvious morphological substructures (Fig. 2A, B). This organization of the SU neuropil into several subdomains was further supported by the combinatorial expression pattern of various cell adhesion molecules. For example, we found the synaptic cell adhesion molecule Teneurin-m to be broadly expressed throughout the AOTU neuropil with the exception of the central subdomain of the SU (SU-c) and the anterior half of the SU-l (Fig. 2C). On the other hand, the adhesion molecules Connectin and Capricious were specifically expressed in the SU-c and SU-m domains, respectively (Fig. 2D, E, F, G). We then tested whether the SU subdomains matched different classes of MeTu afferents (Fig. 2C-K). Based on the terminal arborization patterns from 13 independent expression lines (see material and methods) we could distinguish at least three distinct, non-overlapping populations of MeTu neurons. Based on the segregation of their axons within the AOTU, these neurons were classified as MeTu-lateral (-l), MeTu-central (-c) (partly analogous to the intermediate subdomain from (Omoto, Keles et al. 2017), see below), and MeTu-medial (-m) (compare Fig. 2N-O). A more detailed analysis of molecular markers in combination with MeTu expression lines revealed a further subdivision of the lateral SU domain (SU-l) into distinct anterior and posterior subdomains (SU-l_a_ versus SU-l_p_, Fig. 2C’, F’), which was not apparent for the LU (Fig2 C’, D’, E’ G’). Furthermore, by combining independent Gal4 and LexA expression lines, a similar anterior-posterior division of the central SU domain (SU-c) into SU-c_a_ and SU-c_p_ subdomains was found (Fig. 2H). Importantly, the terminals of specific MeTu driver lines co-labeled specifically with neuropil markers defining these specific subdomains of the SU, indicating that specific subdomains are indeed targeted by specific MeTu classes (Fig. 2J, H’). In contrast other expression lines did not manifest such intra-domain restriction (Fig. 2K).

**Figure 2:**
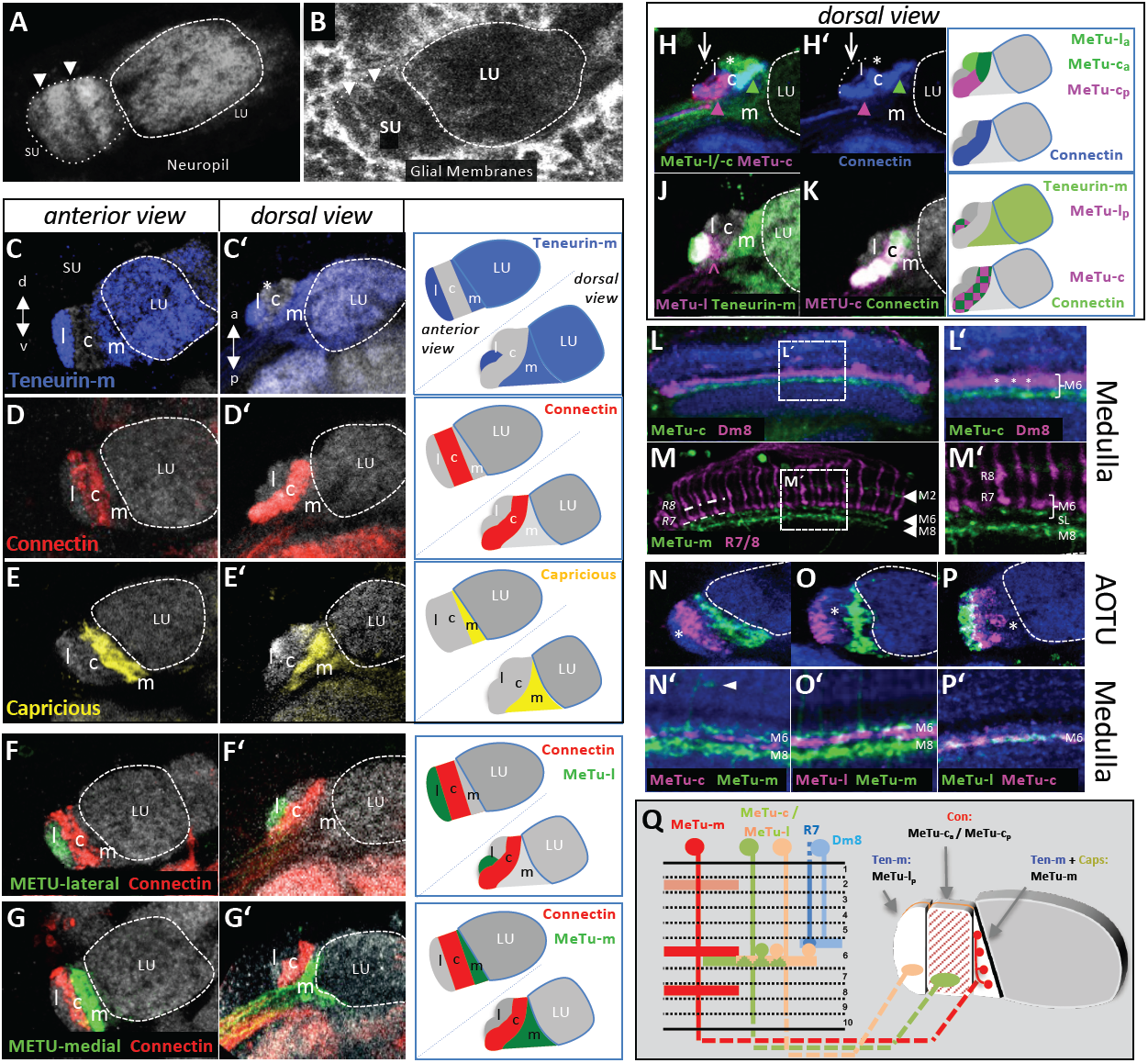
Classification of METU neuron subtypes. **A.** Subdivision of AOTU’s small unit (SU) can readily be observed with neuropil markers (anti-CadN). Arrowheads indicate borders of subdomains. In contrast, the large unit (LU) has a uniform appearance. **B.** Glial labeling using repo-Gal4 reflects the compartmentalization of the AOTU’s SU (arrowheads). **C-E.** Each SU domain is characterized by a unique combination of three cell-adhesion molecules: Teneurin-m (blue) is strongly expressed in the lateral domain (C), with lower intensity in the medial domain and the LU. The lateral domain is further divided into an anterior, Ten-m negative (asterisk) and a posterior, Ten-m-positive compartment (C’). Connectin expression (red) defines the central domain (D, D’). Caprecious-Gal4 (yellow) marks the medial domain (E, E’). **F, G.** Domain borders are respected by terminals of METU subtypes: different Gal4-labeled METUs innervate either the lateral (F-F’) or medial domain (G-G’), without overlapping into the central, Connectin-positive (red) domain. **H-K.** Further division of the lateral and central domain into anterior and posterior compartments: Combination of LexA- (green) and a Gal4- (magenta) lines revealing a subdivision of the central domain (H). Anti-Connectin (blue) labels the complete central domain (H’). A small subset of LexA-expressing neurons also innervates the anterior part of the lateral domain (asterisk). A population of METU-l neurons innervating exclusively the posterior part of the lateral domain, which is also defined by Teneurin-m expression (green). An asterisk marks the anterior, unlabeled part of the lateral domain. The arrowhead marks turning axons not part of the target field. The complete central, Connectin-positive (green), domain is labeled by an line specific for METU-c neuros (magenta). **L, M.** Dendrites of METU-c neurons (green) are restricted below medulla layer M6, in a sublayer below R7 terminals and Dm8 neurons (magenta). Three medulla layers are occupied by METU-m (arrowheads). Photoreceptors are labeled with anti-Chaoptin (24B10). SL, serpentine layer. **N-P.** METU-c/l neurons and METU-m neurons do not overlap in the medulla (N’-P’). Asterisks indicate the respective unlabeled sublayer. In the upper row (N, O, P) the AOTU expression of the same specimen is shown. METU-c and METU-l terminals are separated in the SU, while sharing the same medulla layer. Arrowhead in N’ points to MeTu-m dendrites in M2. **Q.** Schematic overview over METU neuron subtype morphology in medulla and SU.

To get further insights into the neuronal identity of the different MeTu populations, we visualized their dendritic arborizations in the medulla neuropil (Fig. 2L-P’). Interestingly, all three MeTu classes formed dendrites below medulla layer M6, the region where UV-sensitive R7 photoreceptors target their main synaptic partner, the distal medulla cell type Dm8 (Gao, Takemura et al. 2008, Karuppudurai, Lin et al. 2014, Ting, McQueen et al. 2014, Nern, Pfeiffer et al. 2015). MeTu dendrites therefore segregated from the R7/Dm8 synaptic area (Fig. 2 L, M). For the majority of MeTu-l and MeTu–c neurons, the area below layer M6 appeared to be the only layer with dendritic signal (Fig. 2L, P’). In contrast, MeTu-m neurons formed dendritic arborizations in two medulla layers located both proximal and distal to layer M6, most likely layer M2 and layer M8 (Fig. 2M, M’, N’, O’). Interestingly, below medulla layer M6, MeTu-m dendrites segregated from MeTu-l/-c dendrites (Fig. 2N’,O’), thereby revealing three distinct sub-layers within this medulla layer (R7/Dm8, MeTu-m, MeTu-l/-c) (Fig. 2Q). In summary, the AOTU receives direct input from distinct types of MeTu neurons, which differ in their dendritic position, target subdomain, and molecular identity (summarized in Fig. 2Q).

### Photoreceptor connectivity of topographic MeTu subtypes

To investigate whether direct synaptic contacts between MeTu-l/-c dendrites below layer M6 and inner photoreceptors R7 (and less likely R8) might exist, the trans-synaptic tracer ‘trans-Tango’ (Talay, Richman et al. 2017) was used. TransTango was expressed under the control of either R7- or R8 specific Rhodopsin Gal4-driver combinations, respectively (see material & methods; Fig. 3A,B). Significant labeling of the SU was detected following the transTango expression in R7 (A’), whereas no signal was detected in the AOTU in the case of R8 > transTango (B’). In the former case, the obtained patchy signal indicated that only UV-sensitive R7 cells are indeed synaptically connected to some, but probably not all MeTu-l/-c neurons. Although dendrites of MeTu-l and MeTu-c cells were mostly restricted below medulla layer M6, we noticed that some MeTu cell clones formed vertical processes reaching beyond medulla layer M6 (almost reaching M3), thereby making R7 photoreceptor → MeTu synapses a possibility (see MeTu-l clone in Fig. 3C). In order to systematically test which MeTu subtypes are post-synaptic to R7 photoreceptors, we generated a transcriptional fusion of a ∼3.5 kb fragment containing the promoter sequences of the histamine receptor Ort, driving expression of membrane tagged mCD8:GFP (see materials and methods). Since Histamine is the neurotransmitter expressed by all insect photoreceptors (Stuart 1999), their synaptic targets can be marked by ort expression (Gao, Takemura et al. 2008). As expected, this ort-mCD8:GFP transgene labeled many cell types throughout the optic lobes (Supplemental data), as well as MeTu axon projections into discrete domains of the AOTU (Fig. 3D). Out of the five domains of the SU, only three were clearly positive for ort-mCD8:GFP, namely SU-l_a_, SU-c_a_, SU-c_p_. We therefore proceeded to confirm that processes from MeTu subtypes terminating in these domains indeed co-labeled with GFP, using a combination of different subdomain-specific drivers. Out of both MeTu-l subtypes, only axons of MeTu-l_a_ neurons co-labeled with GFP, whereas MeTu-l_p_ did not (Fig. 3E,F). In contrast, axons from both MeTu-c subtypes (c_a_ and c_p_; both individually labeled using different driver lines), co-labeled with GFP (Fig. 3G-I). Finally, axons of MeTu-m cells never co-labeled with GFP (Fig. 3J). In summary, of all MeTu cells innervating the SU of the AOTU, only MeTu-l_a_, only MeTu-c_a_, and MeTu-c_p_ were identified as direct synaptic targets of R7 photoreceptors (Fig 3K).

**Figure 3:**
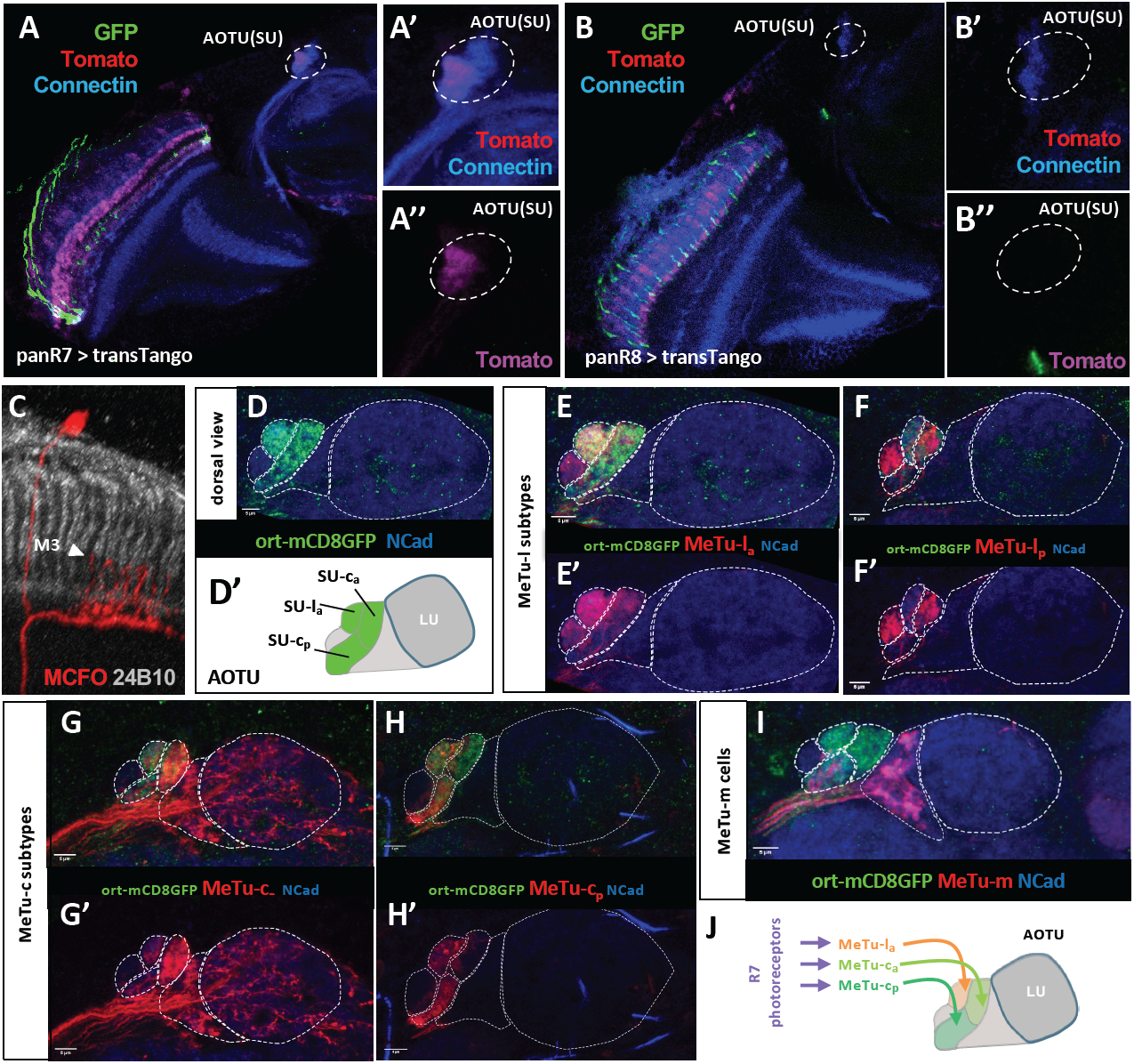
Connectivity between photoreceptors and METU neurons. **A.** R7-specific trans-tango experiment using (rh3+rh4)-Gal4 (‘panR7) reveals tomato-positive transTango signal in METU processes to the SU of the AOTU (dashed area in A’ and A’’). **B.** No transTango signal is detectable in (rh5+rh6 / ‘panR8’) > transTango experiments (B’ and B’’). **C.** Single cell METU-l clone visualized via R52H03 > MCFO-1 reveals an exemplary neuron with dendrites in multiple medulla layers and processes reaching to higher medulla levels (arrowhead in layer M3). **D.** Expression of the newly generated ort-mCD8:GFP transgene in the AOTU. The domains of the SU are labeled (SU-l_a_, SU-c_a_, SU-c_p_), whereas the LU is not labeled (D’). **E.** METU-l driver R94G05 labels both METU-l_a_ and METU-l_p_ populations, yet only METU-l_a_ are post-synaptic to photoreceptors (E’). **F.** METU-l driver R52H03 specifically labels METU-l_p_ and METU-c_a_ populations, of which only METU-c_a_ are post-synaptic to photoreceptors (F’). **G.** METU-c driver R67C09 specifically labels METU-c_a_ cells, which are post-synaptic to photoreceptors (G’). **H.** METU-c driver R25H10 specifically labels METU-l_a_ and METU-c_p_ populations, both of which are post-synaptic to photoreceptors (H’). **I.** METU-l driver R20B05 labels METU-m cells, which are not post-synaptic to photoreceptors (I’). **J.** Schematic summary of the results from D-I. For genotypes, see supplemental Data.

### Topographic organization of AOTU afferents

Next, we proceeded to a more systematic characterization of how AOTU subdomains correlate with MeTu-neuron identity at a single cell level. Clonal analysis revealed a stereotypical, subtype-specific pattern of MeTu innervation, where any given MeTu axon terminates in only one of the five SU subdomains (Fig. 4A-C). For MeTu-l and MeTu-c neurons, a spatially restricted termination pattern was observed in their respective SU subdomains (Fig. 3A, B). In contrast, afferent arborizations of METU-m cells extended throughout a large portion of their compartment (Fig. 3C), resembling the previously published projection pattern of LC10 neurons in the LU (see Fig. 1J). The differences between MeTu-m neurons (with dendritic arborizations in multiple medulla layers and axonal convergence throughout their SU subdomain) versus MeTu-l + MeTu-c neurons (with dendrites restricted below medulla layer M6 and spatially restricted axon terminals in the AOTU) therefore support the existence of morphologically and functionally distinct visual channels into the central brain.

**Figure 4:**
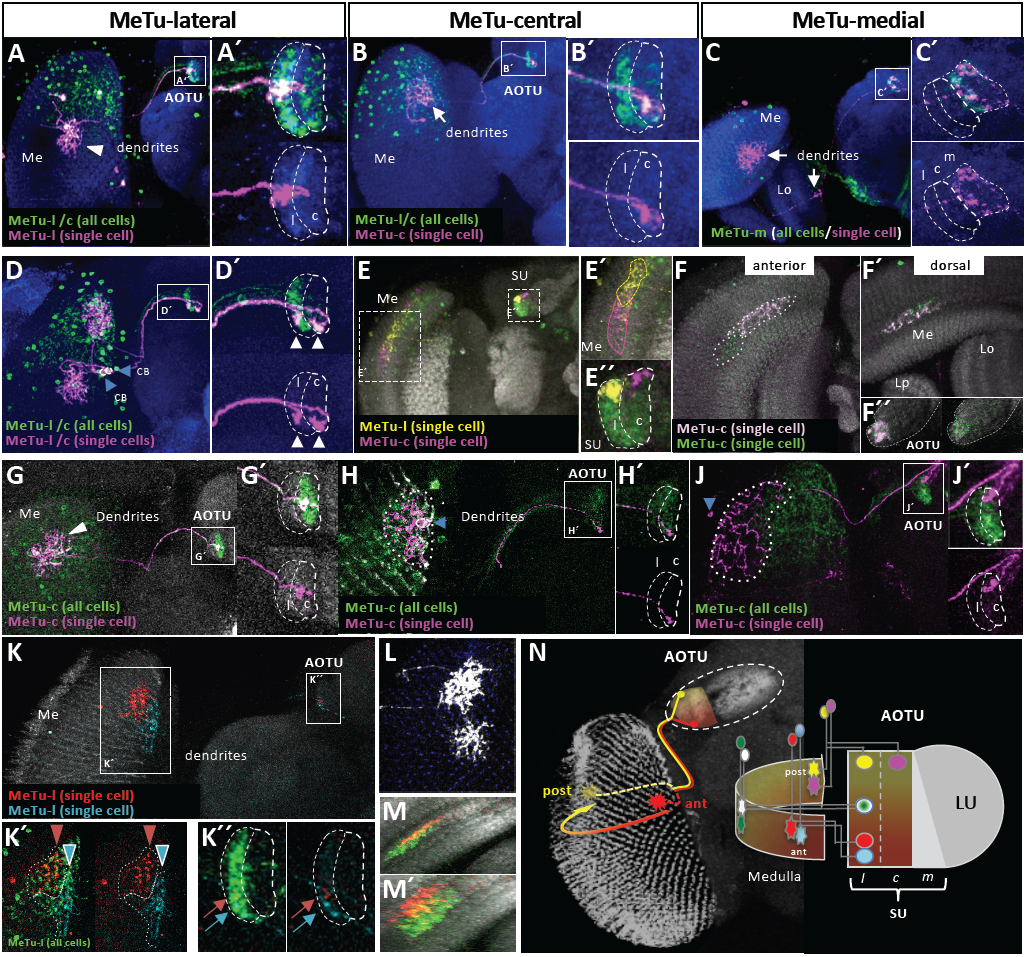
Topographic organization of AOTU projections. **A-C.** FLYBOW-labeling of METU-neurons innervating their respective domain of the SU (magnified in A’, B’, C’). Arrow in (C) indicates innervation of the lobula by METU-m neurons. **D.** Two neighboring cells (blue arrowheads) innervate different positions within the dorsal medulla and targeting to the lateral or the central SU-domain, respectively (white arrowheads). CB, cell body. **E.** Two METU clones with overlapping dendritic fields at the posterior edge of the medulla target to the dorsal edge of either the lateral domain (yellow neuron) or the central domain (magenta neuron), respectively. **F.** Anterior-posterior, but not dorso-ventral positions in the medulla correlate with topographic projections in the AOTU: METU-c neurons at the same A-P position in the medulla target into an overlapping region in the central SU-domain. (Different angles of the same brain are shown). **G-J.** Topographic projections of METU-c neurons: Central dendritic fields in the medulla correlate with central termination the AOTU (G’), anterior dendritic positions in the medulla correlate with ventral targeting (H’), while posterior medullar dendrites correlate with dorsal termination (J’). Size of dendritic fields and size of innervated target area did not correlate (blue arrowheads indicate cell bodies). **K.** Dendritic fields of neighboring clones at the anterior rim of the medulla maintain their topography in the AOTU: The red clone, being located more posteriorly in the medulla terminates at a more dorsal position in the AOTU. **L.** The size of dendritic fields varies amongst METU-l neurons. **M.** Overlap of dendritic fields between two METU-l clones (different angles of the same brain are shown). **N.** Summary of the FLYBOW-pairs described above (colors accordingly) and model of topographic relationships between medulla dendritic fields and SU axis of innervation. For genotypes, see supplemental Data.

Dendritic fields of single METU-neurons always covered multiple medulla columns, yet the specific field size of individual METU-neuron clones varied substantially, both within and between classes (Fig. 3L). Importantly, the differential labeling of randomly induced two-cell clones for either MeTu-l and MeTu-c neurons (using FLYBOW (Hadjieconomou, Rotkopf et al. 2011), see materials & methods) manifested two crucial features with regard to the spatial organization of their terminals in the AOTU: First, MeTu-neurons of the same type (l/l or c/c) with neighboring dendritic fields in the medulla always projected to adjacent positions in the corresponding SU domain (Fig. 4K). Secondly, MeTu-neurons of different types (l/c) with overlapping dendritic fields in the medulla always projected to the same position along the d-v axis, yet in in adjacent SU domains (Fig. 4E). To determine whether MeTu-l and MeTu-c cells innervated their corresponding SU domain in a topographic fashion, we correlated their relative position of dendrites in the medulla with their axon terminals and AOTU, respectively (Fig. 4G-J). For both cell types we could observe a strict correlation between the dendritic position along the anterior-posterior (a-p) axis in the medulla and the axonal termination point along dorso-ventral (d-v) axis in the AOTU (Fig. 4N, n=35). According to this wiring scheme, MeTu-l and MeTu-c neurons with dendrites at the anterior rim of the medulla neuropil target the most ventral position in their corresponding SU domain whereas neurons with dendrites at the posterior rim of the medulla connect to a dorsal edge of the SU (Fig. 4H, J). Furthermore, MeTu-(l/c) clones with dendrites in more medial medulla regions also targeted to medial position in the AOTU (Fig. 4K). In contrast, the spatial arrangement of METU dendrites along the d-v axis of the medulla was not reflected by a distinct targeting pattern the SU domains (Fig. 4F). As expected, no obvious spatial correlation between dendrite position targeting could be detected for METU-m neurons, since their terminals were spreading the entire target domain (not shown). In summary, these data revealed the structural organization of a retinotopic representation in the AOTU part of the central brain of *Drosophila* in which a topographic information (MeTu-l and MeTu-c) is conveyed separately from non-topographic channel (MeTu-m), thereby forming multiple parallel channels.

### AOTU efferents maintain domain identity and visual topography

If the AOTU served as a relay station of spatial information from the optic lobes to central integration centers of the brain, one would expect a matching pattern of connections between MeTu subtypes and corresponding AOTU output neurons along the d-v axis, at least for the lateral and central SU domains. We identified a large set of expression lines for AOTU projection neurons targeting the bulb region (TuBu neurons) (Omoto, Keles et al. 2017). These TuBu expression lines show domain-specific restriction of their dendritic fields, corresponding to the SU-l, -c and -m domains and were therefore classified as TuBu-l, -c & -m neurons, respectively (Fig. 5B, G; compare also (Omoto, Keles et al. 2017)). In single cell clones, the size of the dendritic fields of TuBu neurons matched the extent of axonal arborizations from corresponding MeTu cells. In agreement with subdomain-specific connectivity, TuBu -l and -c domains manifested the most restricted dendritic arbors whereas TuBu-m formed broad dendritic fields (Fig. 5H, J). In addition, the average number of TuBu neurons covering a given SU domain along the d-v axis corresponded well to the average number of MeTu neurons representing the a-p axis in the medulla (between 8-12 neurons for different classes), allowing for a topographic one-to-one representation. To test if the spatial overlap of MeTu axon terminals and TuBu dendrites was indicative of synaptic connections we used the activity dependent GRASP technique (Karuppudurai, Lin et al. 2014, Macpherson, Zaharieva et al. 2015). Indeed, GRASP between presynaptic MeTu neuron subtypes and various sets of TuBu neurons revealed a strict matching of synaptic partners within, but not across SU domains (Fig. 5C-D).

**Figure 5:**
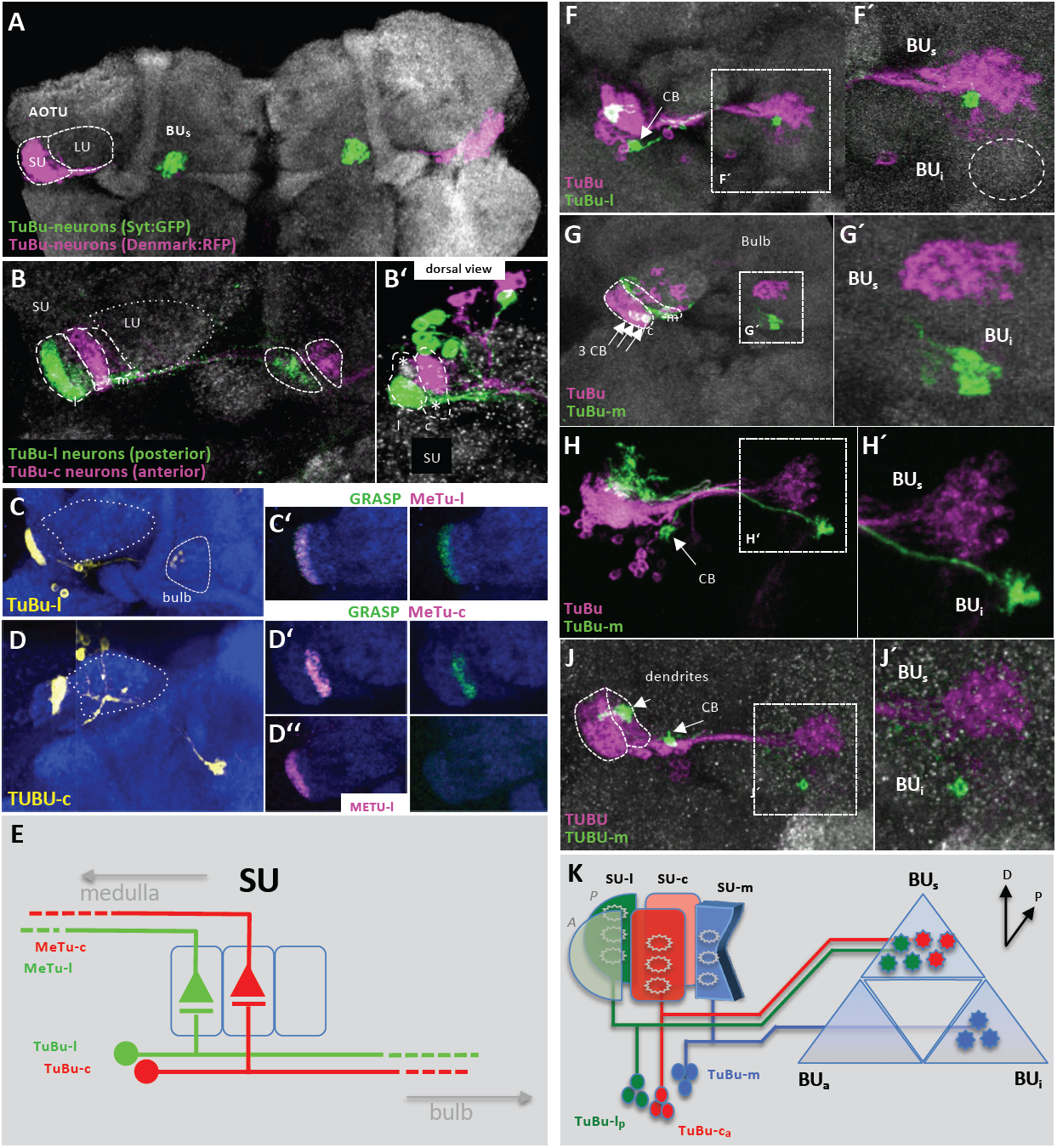
Bulb-innervating neurons descending from the AOTU maintain domain identity. **A.** The bulb neuropil receives input from all three SU-domains. **B.** Terminals of TUBU-l and TUBU-c neurons are spatially separated within the bulb (asterisks mark the unlabeled anterior-lateral and posterior-central SU-domains). **C, D.** Pre-to postsynaptic matching of domain-specific expression lines in the SU revealed by synGRASP: Anti-GFP (yellow) detects the presynaptic moiety of TUBU-l, expressed under Gal4-control. Positive GRASP-signal is obtained in combination with METU-l neurons (C’). **D.** TUBU-c neurons (yellow) are synaptic partners of METU-c neurons (D’), whereas no synaptic connections are formed with METU-l neurons (D’’). **E.** Scheme depicting how afferent METU neurons and efferent TUBU neuron subtypes form circuits in their respective SU-domains. **F-J.** FLYBOW-labeling using a reporter for the majority of TUBU neurons. TUBU innervations are virtually absent from the BUi (dashed circle). CB, cell body. TUBU-l dendrites and axonal terminals are spatially restricted (F’). Three TUBU-m clones innervate a ventral area in the bulb (BUi), separate from TUBU-l&c neurons (G’). TUBU-m arborization size is variable both in AOTU and bulb, ranging from covering larger areas (H) to spatially restricted (J). **K.** Schematic describing the distribution of three TUBU classes in the bulb neuropil. For genotypes, see supplemental Data.

### Dynamic organization of AOTU efferents in the bulb region

TuBu axons form a single fascicle which extends from the AOTU towards the bulb, where they then segregate towards distinct domains according to their SU domain identity (Fig. 5K; compare also (Omoto, Keles et al. 2017)): We found that TuBu-l and –c neurons terminated in adjacent regions of the superior bulb (BU_s_), whereas axons of TuBu-m neurons targeted into the inferior bulb (BU_i_) (Fig. 5F, G). Hence topographic and non-topographic visual pathways remain spatially segregated within the bulb. We next analyzed the spatial organization of dendritic and axonal arborization of single cell and small size TuBu clones. To determine if the retinotopic representation from the AOTU is translated into the terminals of TuBu cells within the bulb region, we compared the relative positions of TuBu dendrites in the SU with the location of their axon terminals in the bulb region by generating two-cell clones within a population of TuBu-l and TuBu-c neurons, respectively (Fig. 6A-C). This analysis revealed that adjacent dendritic positions in the AOTU are indeed maintained within neighboring domains of the bulb, although their relative position to each other within the bulb area is variable (Fig. 6A, B). To further characterize the spatial patterning of TuBu neurons we generated a series of single cell clones and compared the relative position of TuBu dendrites in the SU with their axon termination areas in the bulb, this time for individual TuBu clones (Fig. 6D). In contrast to the strict spatial correlation between MeTu neuron dendrite position along the a-p axis and its axon termination along the d-v axis, the position of TuBu dendritic fields within the SU domain did not predict their site of axon termination within the bulb area (Fig. 6E, F). For example, single TuBu-l clones with dendritic fields in the dorsal SU domain manifested projections either to the dorsal, ventro-lateral, or ventral-medial bulb domains (Fig. 6F, left column). Similarly, the dorsal bulb region could receive TuBu afferents from neurons with either dorsal, medial, or ventral SU positions (Fig. 6F, right column). Given the fixed spatial proximity of TuBu axon terminals with adjacent dendritic fields described above, these data suggest that the topographic map of the AOTU is maintained in the bulb were it translates into a more dynamic organization regarding the a-p and d-v axes of TuBu terminals within a sector of the bulb.

**Figure 6:**
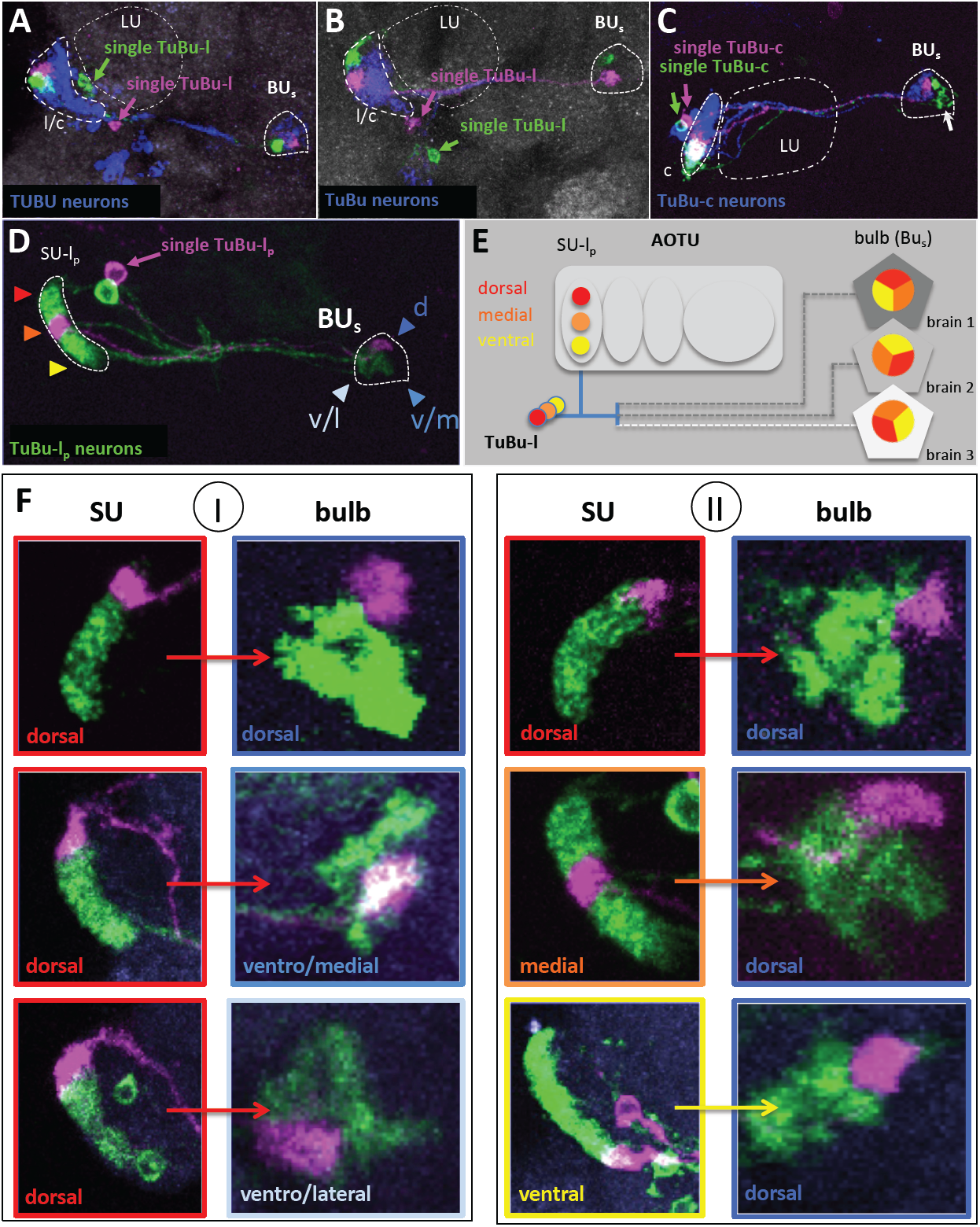
Variability of innervation patterns across TUBU neurons. **A-C.** The axon terminals of neighboring TUBU-l neurons maintain their proximity in the bulb, but their orientation is variable, both when labeling all TUBU neurons (A,B), or TUBU-c specifically (C). **D.** Example of a FLYBOW-induced TUBU-l single cell clone (magenta), while co-labeling all TUBU-l_p_ neurons (green). Color coding of arrowheads indicates dorso-ventral distribution in the AOTU as well as positions in the BUs (dorsal, ventro-lateral, ventro-medial), same as in subsequent panels. **E.** Schematic depicting the lack of stereotypic orientation of terminals from adjacent TuBu-l_p_ neurons in the bulb. **F.** There is no topographic correlation between dendritic position in the AOTU and target field in the bulb. Neurons with dorsal positions in the AOTU target to various positions within the lateral sector of the BUs (column I). Likewise, a similar position in the bulb are innervated from various positions along the D-V axis in the AOTU (column II). For genotypes, see supplemental Data.

### Projections of AOTU domain identity onto ring neurons of the EB

Efferent neurons from the bulb region have been shown to target specific ring layers within the EB (R neurons) (Wolff, Iyer et al. 2015, Franconville, Beron et al. 2018). To characterize the matching between TuBu cells and the spatial positioning of R neuron subtype dendrites, we performed a series of co-labeling studies (Fig. 7 A-F), which, for technical reasons, focused on two TuBu-classes: TuBu-l_p_ & TuBu-c_a_ in combination with different candidate R neuron types of the BU_s_: R2, R4d, and R5. As previously shown, the BU_i_ is innervated by R3 neurons (Fig. 7 G), but not targeted by TuBu-l or TuBu-c neurons (data not shown, compare (Omoto, Keles et al. 2017)). In the BU_s_ we could identify matching projection patterns, in which all TuBu axons of one class appeared to contact only one specific R neuron type. This was particularly clear in the case of TuBu-l cells, which clearly overlap with R4d (Fig. 7A), but not with R2 or R5 (Fig. 7B,C). For TuBu-c neurons, a partial overlap with the dendritic fields of R2 was detected (Fig. 7E), while avoiding contacts with R4d and R5 (Fig. 7D,F). Furthermore, co-labeling revealed that dendrites of different R neuron types segregate into coherent, non-overlapping domains within the bulb neuropil (Fig. 7G-J). In summary, in our analysis of two representative TuBu classes and three candidate R neuron classes innervating the superior bulb (BU_s_), we could dissect one fully matching pair of TuBu → R neuron circuit, as well as another pair with a partial overlap. Thus, yet another synaptic level is added to the parallel visual pathways described here, as distinct AOTU efferents remain separated and contact different EB ring (Fig. 7K).

**Figure 7:**
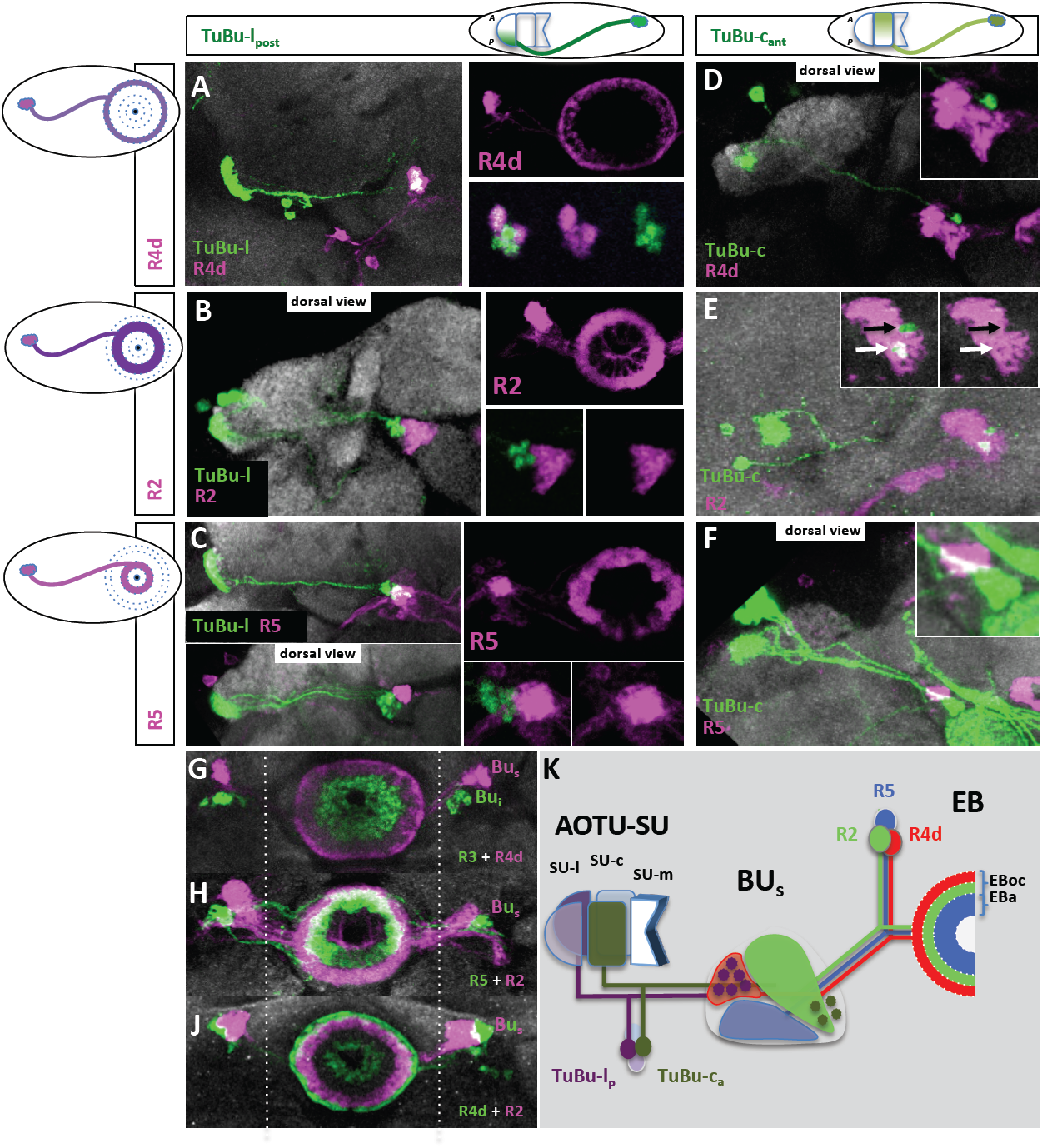
Distinct AOTU pathways connect with specific R neuron classes. In the BU_s_, different TuBu classes connect to a set of R neurons. Two LexA expression lines label the posterior lateral domain and the anterior central domain of the SU, respectively. The BU_a_ and BU_i_ are not covered in this analysis. **A-C)** TuBu-l_p_ neurons (*R25F06-LexA*) innervate the BU_s_, where they exclusively contact R4d neurons, labeled with GM*R12B01-Gal4* (A), but not R2, labeled with EB1-Gal4 (B) or R5, labeled with GMR49B02-Gal4 (C) neurons. **D-F)** TuBu-c_a_ neurons (labeled with *R64F06-LexA*) partially overlap with R2 neurons (E), but not with R4d (D) or R5 (F). White and black arrows in (E) indicate the presence or absence of co-labeling of expression lines, respectively. **G-J)** Co-labeling of R neurons reveals the coverage of different fields within the BU. R3 neurons do not contribute to the BU_i_. Genotypes: (G) *R12B01>mCherry; R14A12-LexA>GFP*, (H) *EB1>mCherry; R48H04-LexA>GFP*, (J) *EB1>mCherry; R85E07-LexA>GFP*. **K)** Proposed segregation of visual information in the superior bulb. Filled dark stars in the BU_s_ indicate terminal endings of TuBu neurons (microglomeruli). EBoc, outer central domain; EBa, anterior domain of EB.

## Discussion

Like various other sensory modalities for which spatial information is critical, neural circuits in the visual system of many animals are organized in a topographic fashion to maintain the neighboring relationship of adjacent pixels detected by photoreceptors in the periphery, along the visual pathways into the central brain (Livingstone and Hubel 1988). The topographic representation of different kinds of sensory information within the central brain of *Drosophila* is currently being investigated using molecular genetic tools in combination with cell-type specific driver lines (Tsubouchi, Yano et al. 2017, Patella and Wilson 2018). Although it is well known that spatially-patterned visual stimuli induce coherent activity bumps in the Drosophila CX (Seelig and Jayaraman 2013, Seelig and Jayaraman 2015, Green, Adachi et al. 2017, Kim, Rouault et al. 2017), the pathway translating peripheral visual information into central activity patterns remains poorly understood.

### Several parallel, topographic pathways into the central brain

Here we have showed that medulla inputs to the AOTU fall into two morphological types regarding their arborization patterns: broadly intermingling terminal axons or spatially-restricted, versus non-overlapping axon terminals. While the topographic representation from the lobula neuropil is mostly lost in the broad arborization pattern of converging LC projection neurons onto the majority of optic glomeruli (Panser, Tirian et al. 2016, Wu, Nern et al. 2016, Keles and Frye 2017), we could identify a unique spatial organization for the output channel from the medulla (Fig. 8). Topographic representation of the medulla (at least its dorsal half, where most driver lines used here are expressed) is maintained in a defined sub-compartment of AOTU (the SU), which is spatially separated from lobula representation within the AOTU (the LU). A strict topographic correlation exists between the a-p position of the dendritic fields of MeTu projection neurons in the medulla and their restricted axon termination along the d-v axis within distinct domains of the SU in the AOTU.

**Figure 8:**
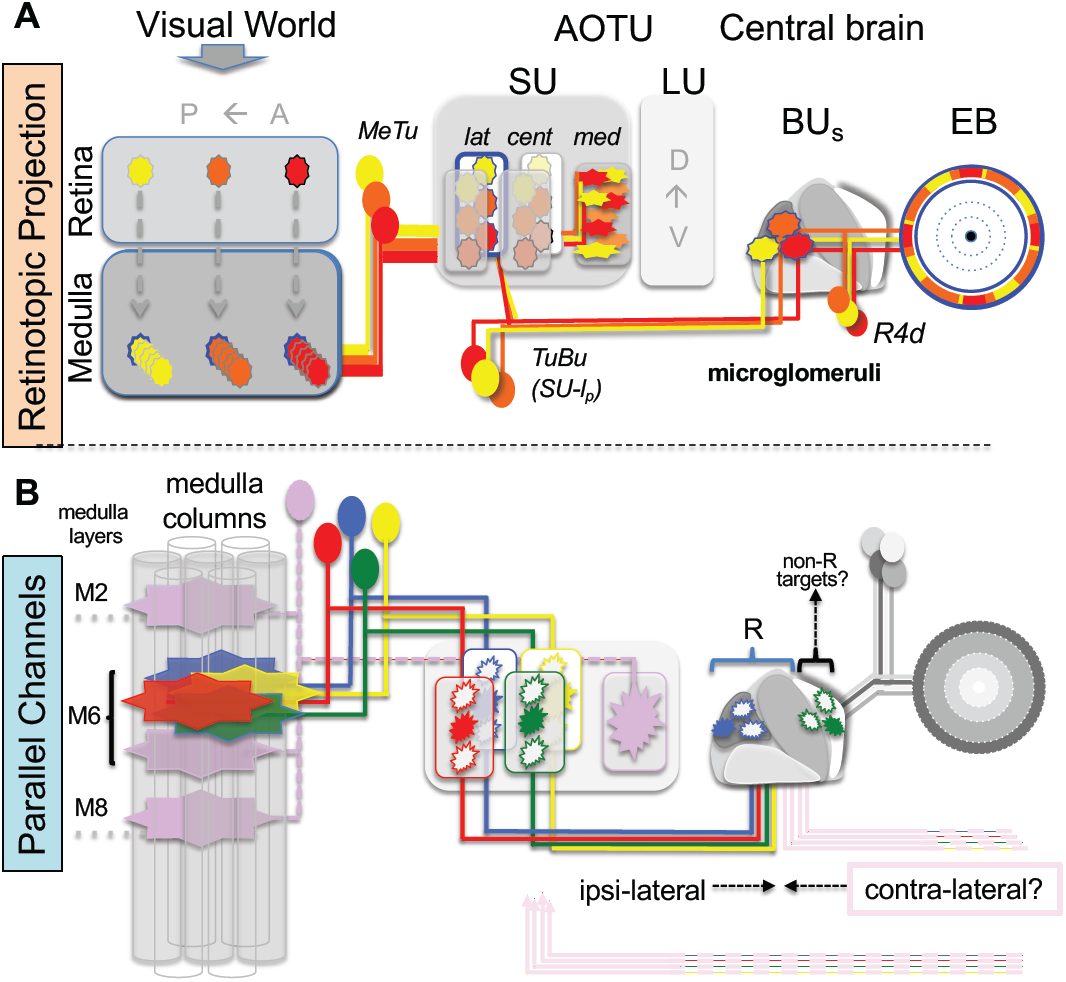
The Anterior Visual Pathway circuit. In the graphic, two features of the pathway - retinotopy and parallel channels - are highlighted. **A.** The retinotopy of the pathway is demonstrated by single neurons. Three spatially separate visual stimuli are transmitted by yellow, orange and red cells, respectively. Innervation patterns in the SU_m_ domain and in the EB indicate a loss of retinotopic arrangements. **B.** Parallel channels exist among several synaptic steps. In the medulla, five neuron classes, innervating separate AOTU compartments, detect visual stimuli from the same medulla columns. For two classes, the target areas in the BUs are shown, where corresponding ring neurons (R) transfer the information into the EB. Inhibitory neurons from the opposite hemisphere are possible regulators in the BU_s_ and the AOTU.

Based on their morphology, as well as their molecular identity, three principle types of MeTu neurons provide input into the AOTU, with overlapping dendritic fields within the medulla but segregated axon terminals to distinct AOTU (sub-)domains. MeTu-l and –c classes have a similar neuronal morphology with dendrite arborization restricted to a single medulla layer (M6) and spatially narrow axon termination areas in four separate AOTU subdomains (l_a_, l_p_, c_a_, and c_p_), thereby building the several pathways arranged in parallel (Fig. 8). Our nomenclature of the SU subdomain organization differs slightly from previous studies (Omoto, Keles et al. 2017), since it is now based on the expression patterns of different cell surface molecules, which we believe reflects the functional organization of these structures. Because of this new classification, both lateral and central domains (but not the medial domain) of the SU become further subdivided into either anterior (SU-l_a_ and SU-c_p_) and posterior halves (SU-l_p_ and SU-c_p_). Nevertheless, it should be noted that the total number of subdomains is not affected. The organization of the MeTu-l and MeTu–c neurons therefore enables a point-to-point projection of visual information that is linked to the columnar organization in the medulla making the corresponding AOTU domains well suited to relay topographic information towards the central brain.

### The transformation of topographic information in the central brain

The borders of the SU compartments are respected by molecularly defined populations of TuBu neurons, thereby defining the next synaptic elements in the parallel pathways towards the bulb neuropil. While this neuropil with its afferent (TuBu) and efferent (R neurons) channels has been intensively studied recently (Seelig and Jayaraman 2013, Seelig and Jayaraman 2015, Green, Adachi et al. 2017, Omoto, Keles et al. 2017, Shiozaki and Kazama 2017, Sun, Nern et al. 2017, Franconville, Beron et al. 2018), there still remains a gap in knowledge concerning how precise synaptic connections convey topographic information to the central complex. Four major findings of the TuBu→EB circuit are revealed by our study: First, the topographic position of TuBu dendrites in the SU is not translated into a defined position within the bulb, but instead exhibits a targeting plasticity within a restricted bulb area. Secondly, while the recent dissection of the AOTU→EB pathways described the bulb as a tripartite structure (Omoto, Keles et al. 2017) including both afferent and efferent neurons, we can now refine this picture by highlighting that, although our analysis of TuBu-neurons is mainly restricted to only two representative TuBu classes (one in the SU-l_p_ and the other in the SU-c_a_ domain), both these classes target to areas within the superior bulb (BU_s_). More broadly expressed driver lines revealed exclusive TuBu neuron innervation of the BU_s_, indicating that additional TuBu classes target to this bulb area (data not shown). Thus, we expect at least four different classes of TuBu neurons to exclusively innervate the BU_s_ (TuBu-l_a_, TuBu-l_p_, TuBu-c_a_, and TuBu-c_p_), each of them connecting to a different set of output neurons, indicating an even more complex organization of the bulb, in particular the BU_s_. Thirdly, TuBu classes project onto dendritic areas of R neuron classes (so called ‘sectors’) within the bulb, and specific connections are formed between TuBu neurons and R neuron classes. Although we could identify three R neuron classes within the BU_s_, there probably exists a much higher diversity of connections within this small area of the bulb, reaching beyond the scope of this study. For instance, the postsynaptic partners of one subset of TuBu-c_a_ neurons as well as neurons contacted by R2 and R5 dendrites remain to be identified. Additional post-synaptic partners other than R neurons are contacted by TuBu neurons, like contralaterally projecting neurons described in the locust (el Jundi and Homberg 2012) and the bumblebee (Pfeiffer and Kinoshita 2012), which connect the AOTU units of both hemispheres (TuTu neurons). Since contralateral inhibition is a known feature of bulb neurons (Sun, Nern et al. 2017) and there remain large, uncovered sectors in the BU_s_, we therefore speculate that projections of contralateral inhibitory neurons might be part of the bulb network described here.

It appears therefore that topography is conserved within the AOTU output neuron projections towards the bulb and ring neurons. All ring neurons of the same type occupy the same ring layer within the ellipsoid body, raising the question of how topographic information is integrated within central complex neuropils. Interestingly, while different AOTU domains receive retinotopic input from the same point in space and are mapped onto TuBu neurons, these parallel channels diverge within the bulb regions, where we found SU-l_p_ and SU-c_a_ efferents mapping onto separate ring neurons (R4d versus R2). Hence we could define at least two distinct topographic MeTu channels into the central brain. While functional differences between the BU_i_ and BU_s_ have been described (Omoto, Keles et al. 2017), functional studies (Seelig and Jayaraman 2013, Sun, Nern et al. 2017) have not yet addressed the neuronal activity of different TuBu classes, or the responses of R neurons within the BU_s_. Based on the data presented here, we would expect that retinotopic information in the BU_s_ remains represented in the respective sector that is associated with their TuBu class.

### An additional, non-topographic pathway into the central brain

A morphologically distinct class of MeTu cells is formed by MeTu-m cells which arborize broadly in their respective AOTU domain. Since the axons of single MeTu-m neurons converge and intermingle within their AOTU domain, which is reminiscent of the afferent organization of LC neurons from the lobula within optic glomeruli in the PVLP regions as well as in the AOTU’s large unit (LU), these cells are well suited to form a non-topographic channel to the central brain. Hence, the AOTU also relays a non-topographic channel (via MeTu-m and TuBu-m cells), in which multi-columnar medullar projection neurons project onto a separate domain, which becomes represented in the inferior bulb area (Bu_i_). Interestingly, while topographic MeTu-l and -c neurons form dendritic fields within a single medulla layer, the non-topographic MeTu-m neurons integrate from three different medulla layers, reminiscent of lobular LC neurons, the main afferents of the AOTU large unit, for which merely rough topography along the dorso-ventral axis has previously been described (Wu, Nern et al. 2016). Furthermore, only non-topographic MeTu-m neurons form a collateral arborization in the lobula, indicating that this pathway could directly integrate visual information from both the medulla and lobula. Nevertheless, the large differences in termination area within the AOTU observed for different MeTu-m single cell clones indicates that these cells may in fact form an even more heterogeneous population. Taken together, topographic and non-topographic afferents generate an interesting assembly of adjacent domains within the AOTU, from exclusively topographic medulla input in SU-l and SU-c domains, non-topographic medulla (and potentially also lobula-) input in SU-m, and another large area of non-topographic input exclusively from the lobula in the LU (Fig. 8). Thus, we have identified multiple parallel topographic pathways separated from a parallel non-topographic channel.

### Evolutionary conservation of the anterior visual pathway

This principle visual pathway involving the AOTU as a central relay station between medullar/ lobular inputs and the central brain is widely shared among different insect taxa, where homologous structures can be found, e.g. orthopterans (Homberg, Hofer et al. 2003), hymenopterans (Mota, Yamagata et al. 2011) and beetles (Immonen, Dacke et al. 2017). The stimuli conveyed by this ‘anterior visual pathway’ have been addressed in only a few insect species so far. Most prominently, the AOTU has been associated with celestial orientation using polarized skylight in several species (Pfeiffer, Kinoshita et al. 2005) or in chromatic processing (Paulk, Phillips-Portillo et al. 2008, Mota, Gronenberg et al. 2013). Dorsal rim ommatidia harboring polarization-sensitive photoreceptors for polarized light vision are crucial for the sky-compass orientation and exist in most insects analyzed, like locusts (Homberg and Paech 2002, Pfeiffer, Kinoshita et al. 2005), butterflies (Labhart, Baumann et al. 2009, Heinze and Reppert 2011) and honeybees (Held, Berz et al. 2016), as well as flies (Wada 1971, Wada 1974, Wernet, Labhart et al. 2003). However, it remains unknown whether MeTu neurons receive direct or indirect input from the DRA. In addition, processing of chromatic information was also shown to be accomplished via the AOTU in several insects (Mota, Gronenberg et al. 2013, Otsuna, Shinomiya et al. 2014). We have now identified the inputs to this pathway, by identifying direct connections between MeTu cells and the color-vision pathway (represented by the UV-sensitive R7 cell) in medulla layer M6.

We describe a combination of cell-surface molecules that serve as landmarks for different AOTU domains and can be further refined by intersectional strategies in order to reveal additional ‘territories’ and distinct neuronal. For instance, as we did not encounter any TuBu neurons targeting the BU_a_ (compare (Omoto, Keles et al. 2017)), we propose that the SU of the AOTU might consist of additional functional units that so far have not been identified. Furthermore, the molecular markers used here can serve as future tools to reveal the molecular mechanisms that underlie the formation of the LC-optic glomeruli network, across species. Since the fruit fly is among the smallest species for which the AOTU has been characterized, and is believed to be a behavioral generalist, even more sophisticated architectures of the SU-homologue could exist in other insect taxa. On the anatomical and functional level, optic glomeruli share many features with the synaptic neuropil within the antennal lobe, which led to the postulation that the glomerular organization in the protocerebrum (optic glomeruli) and the deutocerebrum (olfactory glomeruli) are in fact homologous structures (Strausfeld 1989, Mu, Ito et al. 2012). Indeed, we found molecular characteristics in the PVLP and AOTU that resemble the combinatorial code of cell-surface proteins in the olfactory system (e.g. expression patterns of Ten-m, Con, Caps and Sema1a in both systems). Future developmental studies of mutant LC and METU neurons will reveal to what extent common mechanisms of glomerular circuit assembly exist in both sensory systems. Although the idea of a serial homology of glomerular organized neural system is far from being resolved, it will be intriguing for further studies to analyze the developmental mechanisms that underlie the circuit formation of these parallel AOTU pathways and optic glomeruli circuits as well as to compare them with known molecular functions during olfactory system maturation.

## Materials and methods

### Fly rearing

Flies were maintained in vials containing standard fly food medium at 25°C unless otherwise mentioned. Canton-S flies were used as a wild type strain.

### Fly stocks

The following driver lines were generated at the Fly Light Gal4-/LexA-Collection (Jenett, Rubin et al. 2012) and obtained from Bloomington Drosophila Stock Center (BDSC). One driver line was obtained from the Vienna Tiles collection (Kvon, Kazmar et al. 2014).

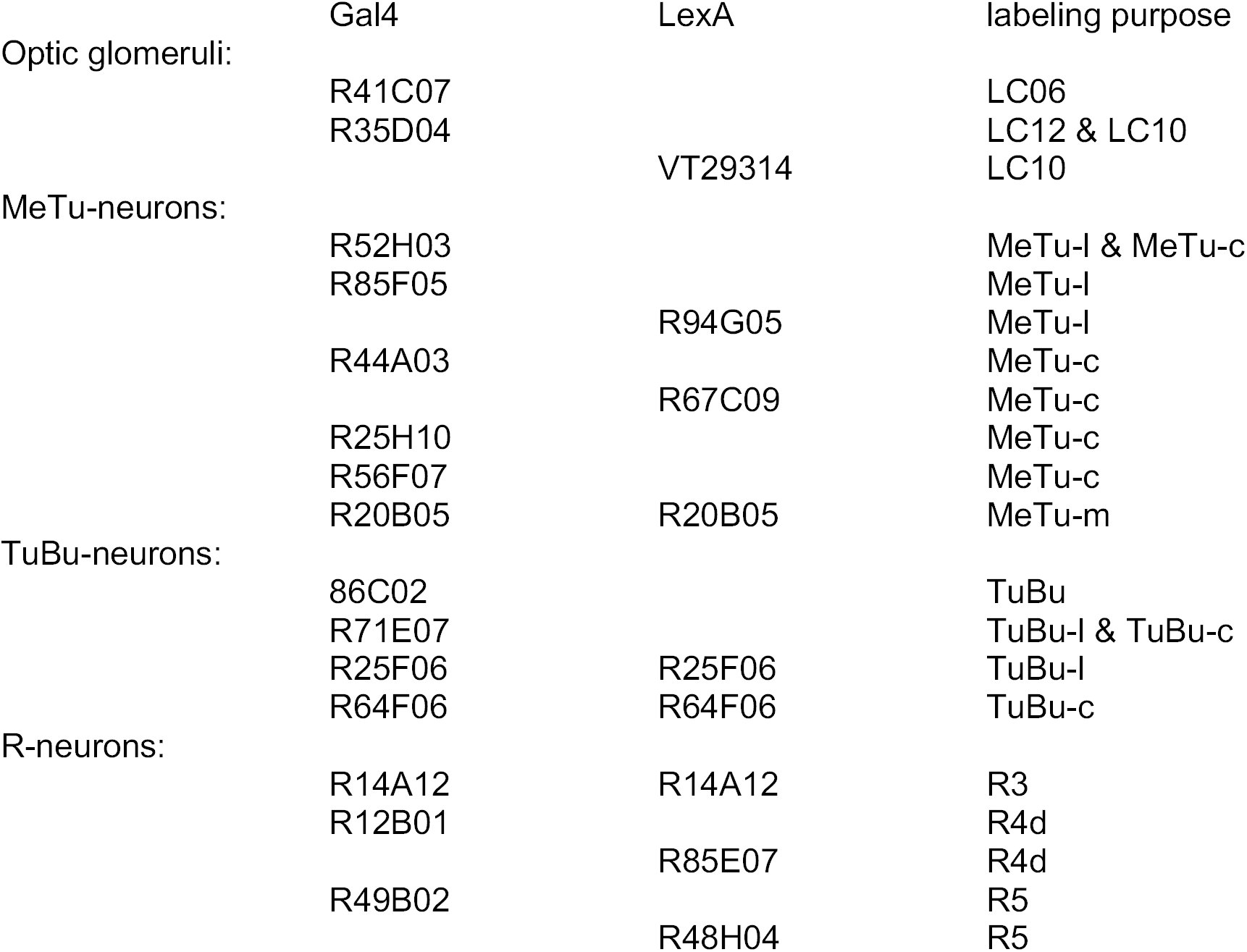

#### Stocks for clonal analysis and effector lines for cell labeling

FRT42D; FRT42D TubP-Gal80; UAS-mCD8::GFP, UAS-mCherry were obtained from BDSC. The UAS-DenMark construct was kindly provided by Bassem Hassan, LexAop::GFP was a kind gift from Andrew Straw. Flies for synaptic-GRASP experiments (UAS-Syb::spGFP1-10 & LexAop spGFP11::CD4) (Karuppudurai, Lin et al. 2014) were a kind gift from Chi-Hon Lee.

#### Generation of ort-mCD8 transgenic flies

A ∼3.5 fragment from the ort gene spanning the entire 5’ intergenic region, as well as the 1^st^ untranslated exon and the transcription start was PCR-amplified, with appropriate restriction endonuclease recognition sites attached to the primers. The fragment was subcloned, sequenced and ligated into a promoterless injection vector (pCasper-mCD8:GFP-SV40). Insertions on 2^nd^ and 3^rd^ chromosomes were obtained via commercial embryo injection. Interestingly, expression was not variegated as seen for many ort-Gal4 constructs. Further information is available upon request.

#### Specific cell labeling

Or67d::GFP and OR67d-Gal4 (Couto et al. 2005) was used for olfactory class visualization, glia cells were marked by repo-Gal4 (Sepp et al. 2001) and Chi-Hon Lee provided the ortC1a-LexA::VP16 (Ting, McQueen et al. 2014) construct for labeling of Dm8 neurons. PanR7-Gal4 (rh3+rh4-Gal4) was used for R7 transTango, and panR8-Gal4 (rh5+rh6-Gal4) for R8 transTango (both gifts from Claude Desplan). Caps-Gal4 (Shinza-Kameda, Takasu et al. 2006).

### Antibodies used in this study

*Primary antibodies* used were: 24B10/Mouse anti-Chaoptin (1:50, DSHB), DN-Ex #8/Rat anti-CadN (1:20, DSHB), Flamingo#74/Mouse anti-flamingo (1:20, DSHB), Rabbit anti-GFP (1:1000, Invitrogen, Carlsbad, CA). Mouse anti-Teneurin-m (1:20) was a kind gift from Stefan Baumgartner, anti-Connectin (1:20) was kindly provided by Robert AH White.

*Secondary antibodies* used: Goat anti-Rabbit Alexa-488 (1:500), Goat anti-Rabbit Alexa-568 (1:300), Goat anti-Mouse Alexa-488 (1:300), Goat anti-Mouse Alexa-647 (1:500), Goat anti-Rat Alexa-647 (1:300). All secondary antibodies were obtained from ThermoFisher Scientific (Alexa Fluor®, Molecular Probes™).

### Clonal analysis

Two approaches for visualization of large and small genetic mosaics were used respectively. For labeling larger mosaics, MARCM clones with an ey-Flp insertion on the X chromosome (Newsome, Asling et al. 2000) were generated as previously described (Lee and Luo 1999). For small clones and single-cell analysis, we used the temperature-sensitive hs-mFlp5 promotor in combination with a FLYBOW (FB1.1B)-construct (Hadjieconomou, Rotkopf et al. 2011, Shimosako, Hadjieconomou et al. 2014). Prior to screening for brains with single cell labeling, a heat shock was given to developing flies (L2-stage, L3-stage, early pupal) for 30min, 1h or 2h at 38°C. The pupae were then allowed to further develop at 25°C and dissected within two days after eclosion.

### Immunohistochemistry

Drosophila brains were dissected in phosphate-buffered saline (PBS) and fixed in 4% paraformaldehyde (PFA) in PBS for 20 min. Samples were washed 3 × 15 min with PBST (PBS containing 0.3 % Triton X-100) and blocked for 3 hours (10% Goat serum in PBST) under constant shaking on a horizontal shaker, before incubating in primary antibody solution for two days at 4°C. Washing procedure was repeated before incubating with secondary antibody for two days at 4°C. Following three times washing, the brains were mounted in Vectashield® (Vector Laboratories, Burlingame, CA) anti-fade mounting medium prior to confocal microscopy. Images were obtained using a TCS SP5II confocal microscope (Leica) using 20x and 63x glycerol immersion objectives. Image processing was performed using ImageJ and Adobe Photoshop® CS6.

### Activity GRASP

Flies were grown in a 25° C, 12h-12h dark-light cycle incubator in normal vials and transferred to custom-made UV-transparent Plexiglas tubes [wall thickness: 4mm] before light induction. 1-day old flies were kept in a 25°C, 20 h – 4 h light-dark cycle custom-made light box (including unpolarized UV light) for 3 days to ensure syfficient photoreceptor activation in the DRA.

Dissection & staining occurred as described above. Brains were stained with polyclonal GFP (anti GFP goat pAB) and monoclonal GFP (anti-GFP rat mAB) antibody to visualize postsynaptic cells and GRASP signal, respectively. Post-synaptic cells were visualized by staining with CD4 antibody.

### Trans-Tango

Flies for trans-Tango experiment were either kept in 18° C or 25° C, in 12h light/dark cycle incubator and dissected when they were either 3, or 15 days old (depending on the experiment).

## Acknowledgements

The authors would like to thank Chi-Hon Lee, Andrew Straw, Tom Clandinin, and Claude Desplan for fly stocks and reagents. This work was supported by the Deutsche Forschungsgemeinschaft (DFG) through grants WE 5761/2-1, SFB958 (Teilprojekt A23), through AFOSR grant FA9550-19-1-7005, through the Berlin Excellency Cluster NeuroCure, with support from the Fachbereich Biologie, Chemie & Pharmazie of the Freie Universität Berlin, as well as the Division of Neurobiology at Freie Universität Berlin (support of FU Berlin and the National Institute of Health to Robin Hiesinger).

## Notes

### Competing Interest Statement

The authors have declared no competing interest.

